# Colicin Receptor CirA Enhances *Salmonella* Typhimurium’s Resistance to Colicin Ib in the Absence of the Cognate Immunity Protein

**DOI:** 10.1101/2024.06.29.601355

**Authors:** BC Gollan, L Luo, Yan Li, J Clark-Corrigall, B Qadri, A Alshuwaier, J Hinton, CMA Khan

**Author notes:** Correspondence: BCG, CMAK.

## Abstract

Intestinal microbiota play a central role in colonisation resistance providing a fundamental barrier to infection to enteric pathogens. An important mechanism of colonisation resistance involves the production of antimicrobial peptides, such as colicins. Pore-forming colicins, synthesised by *Escherichia coli* (*E. coli*) strains, target competing bacteria in their environmental niche, whilst the producing cells are safeguarded by specific immunity proteins. Notably, non-typhoidal *Salmonella* Typhimurium strains can produce a narrow-spectrum protein toxin colicin IB (ColIb) providing a competitive edge against susceptible *Enterobacteriaceae* strains. However, the multi-drug resistant and systemically invasive iNTS (invasive non-Typhoidal *Salmonella*) *S*. Typhimurium D23580 strain poses an interesting case. The strain lacks colicin Ib production and the corresponding immunity protein, but its potential vulnerability in a colicin-rich gastrointestinal milieu remains uninvestigated. In this study, *S*. Typhimurium D23580 exhibited resistance to colicin Ib under tested conditions, despite the absence of the immunity gene *imm*. Intriguingly, in colicin Ib-producing *S*. Typhimurium strains, the *imm* gene appeared functionally redundant in contrast to our current understanding. ColIb binds to the outer membrane protein CirA and is translocated to the inner membrane where it forms a pore in sensitive bacteria dissipating the electrochemical potential. Through a series of experimental approaches, including the use of *Escherichia coli* and *S*. Typhimurium *cirA* deletion mutants, promoter-swap techniques, and gene complementation, we identified that the colicin resistance phenotype in *S*. Typhimurium was partly attributable to the CirA receptor. This finding suggests a complex interplay in the microbial resistance to colicins, highlighting the intricacies of microbial interactions within the gastrointestinal environment.

## Introduction

Colonisation is the initial step for pathogens to establish an infection. Whilst the host deploys several defence mechanisms against enteric pathogens, colonisation resistance plays a pivotal role in preventing colonisation of the host (1). The ability of enteric pathogens to compete and colonise within the high-density population of the intestinal microbiota is key to success. These bacterial enteropathogens have developed numerous strategies to compete with the resident intestinal microbiota (2). The interactions between these pathogens and the intestinal microbiota significantly influence their virulence and pathogenicity, as exemplified by numerous enteric pathogens, including *Salmonella* Typhimurium and Enterohemorrhagic *E. coli* (EHEC) (3, 4).

One mechanism by which bacteria compete within the gastrointestinal tract is through the production of bacteriocins. Bacteriocins are antimicrobial peptides, synthesised by ribosomes, and known for their heat-stability (5). Bacteriocins display a narrow spectrum of activity against closely related strains. While primarily produced by Gram-positive bacteria, certain Gram-negative bacteria, such as *Escherichia coli* also produce bacteriocins. Bacteriocins produced by the *Enterobacteriaceae* family are referred to as colicins (6).

Bacteriocins play a crucial role in colonisation resistance, notably the bacteriocin thuricin is known to inhibit *Clostridium difficile*, and microcins M and H47 produced by *E. coli* have been documented to suppress *S.* Typhimurium (7, 8).

Despite the known presence of numerous bacteriocin producing bacteria within the gastro-intestinal system, the detailed interactions mediated by bacteriocins remain inadequately understood (7, 9). There is growing evidence to suggest bacteriocins play a significant role in the complex interplay between commensal and pathogenic bacteria, this evidence also points to bacteriocins as potential alternatives to traditional antimicrobials. This underscores the importance of exploring how bacteriocins work and their impact on bacterial competition within the gastrointestinal environment. (5, 10–12).

Colicins are typically produced by *Enterobacteriaceae*, and affect closely related bacteria (13). Non-typhoidal *Salmonella* (NTS), specifically *S.* Typhimurium, are known to produce several colicins (14). These proteins are often synthesized under stress conditions, such as the activation of the SOS response system (12). Bacteria that produce colicins are protected from their own toxins through the expression of immunity genes that are co-expressed with the colicin-producing genes (15). Colicins can be categorized into two groups based on their method of entry into target cells. Group A colicins use the Tol system, while Group B colicins enter via the Ton system (15).

Colicin Ib (ColIb) is a Group B colicin encoded by the *cib* gene, and binds to the outer membrane protein CirA, a catecholate siderophore receptor (14). ColIb functions by forming a pore in the inner membrane and disrupting the electrochemical membrane gradient resulting in bacterial cell death (16).

ColIb was first identified in *E. coli*, impacting susceptible members of the *Enterobacteriaceae* family and ColIb has since been demonstrated in several pathogens including *S.* Typhimurium and *Shigella sonnei*, (12, 17, 18). However, ColIb is also produced by *S*. Typhimurium with effects on non-producing *E. coli* strains (6). In contrast to Group A colicins, Group B colicin operons do not encode a gene for cell lysis. It has been demonstrated that the release of ColIb into the environment is facilitated by lysis of cells caused by bacteriophages (19).

The production of colicin Ib offers a competitive advantage to *S*. Typhimurium against commensal *E.coli* during inflammation (14).

Colicins Ia and Ib enter cells by binding to the CirA receptor (15). CirA is a ferric iron-catecholate outer membrane transporter in Gram negative bacteria associated with the uptake of ferric iron under iron limited conditions (20) (21). Zhang (20) showed that CirA played a key role in biofilm formation, motility, antibiotic sensitivity, adhesion and invasion and overall virulence of *S*.

Enteritidis in mice. CirA is a 22 stranded beta barrel with an N-terminal plug domain, where R-domains of colicins colIa/b associate with the ligand binding pocket, where the tip of the colicin R-domain is able to bind to CirA by mimicking siderophore binding (22). In colicin Ia, the binding to the ligand binding pocket of CirA triggers multiple conformational changes, with the most significant changes at the extracellular side, including a substrate induced “opening” of the TonB-dependent transporter by exposure of the plug domain (22).

Colicin I sensitivity is dependent on CirA, where *cirA* deletion mutants are resistant to killing (20). Mutations in genes involved with colicin binding, such as *cirA* and *tonB* are associated with resistance to colicin Ib in the absence of the immunity gene (14, 23–25). ColIb producing strains are protected from the toxic effects of the colicin by expression of the immunity gene (*imm*) which interferes with the interaction with the inner membrane (14). By inserting into the bacterial inner membrane, the immunity protein blocks pore formation and killing by colicin Ib (23). In the absence of this immunity gene, resistance can be acquired through mutations in the colicin receptor, in the case of colicin Ib, CirA. Although the role of CirA as a colicin receptor in the pathogenesis of *Salmonella* is not fully understood is has been suggested that the disruption of CirA, and subsequent colicin resistance, may be a competitive advantage in emerging serotypes (26).

Colicin Ib encoding plasmids harbour operons consisting of the colicin gene (*cib*) and the immunity gene (*imm*), where both genes are tightly regulated in the colicinogenic bacteria. The immunity gene is transcribed in the opposite direction of the pore-forming colicin producing gene (15). In colicin Ib, transcription of *cib* is controlled by both the ferric uptake regulator (Fur) and LexA regulators, as shown in **Figure 1**. LexA is a transcriptional repressor which prevents transcription by binding to a specific binding site. The activation of the SOS operon, such as in the event of DNA damage, results in the autocleavage of LexA, allowing transcription to occur (27).

**Figure 1.**
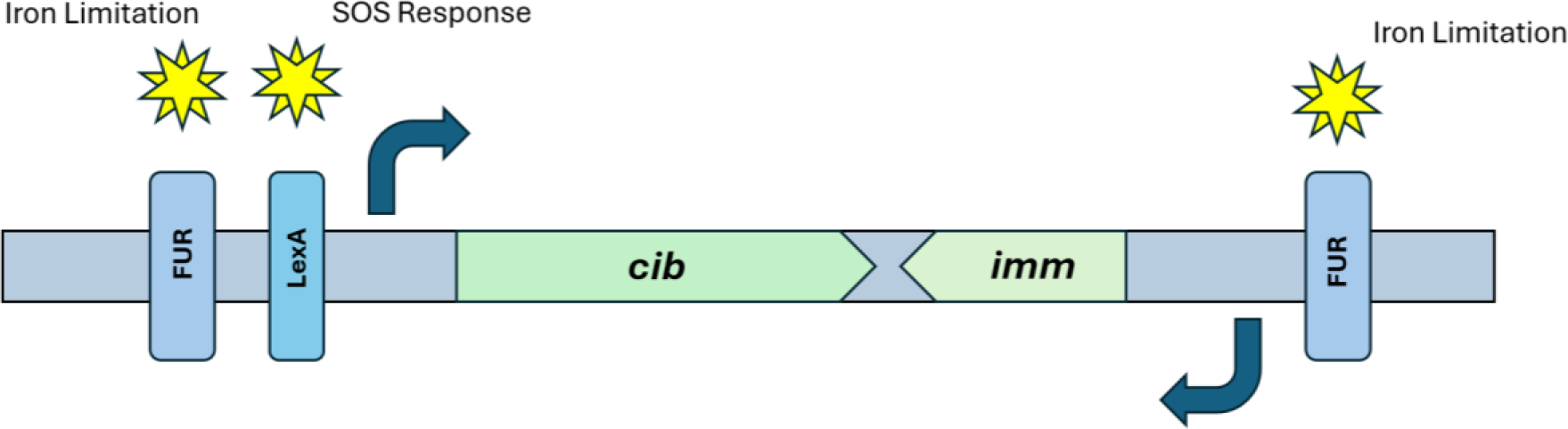
Graphical representation of the regulation of the producing (*cib*) and immunity (*imm*) genes of colicin Ib. Genetic organisation of the colicin Ib locus in *S*. Typhimurium denoting the regulation of the *cib* and *imm* genes by the FUR and LexA transcriptional regulators, adapted from Spriewald (16).

Both the *cib* and *imm* genes are also regulated by the Fur regulator, this iron dependent regulator binds to the ferrous ion Fe^2+^, where in its iron bound state it represses transcription (28). The repression is then released under iron limited conditions. Colicins Ia and Ib are the only colicins to be regulated by both Fur and LexA, where maximal expression requires both stress conditions (stimulation the SOS response) and iron limitation (14).

The competitive benefits of colicin Ib production by *S*. Typhimurium on commensal *E. coli* has been established during inflammation (14). However, our knowledge and understanding into the impact of colicin Ib on systemically invasive non-typhoidal *Salmonella* (iNTS), or of the importance of the immunity gene in colicin-producing strains is limited.

From a genome mining analysis of over 600 microbes and representing almost 200 genera from the human gut, Drissi (29), revealed approximately half the genomes encode for putative bacteriocins. The widespread distribution of bacteriocin genes within the intestinal microbiota suggest bacteriocins may play a role in colonisation resistance of competing pathogens. Determining the effects of colicins, such as colicin Ib on enteric bacteria such as *Salmonella,* is pivotal in understanding the interactions with the intestinal microbiota.

The colicin Ib and immunity protein genes are encoded on the low-copy plasmid pColIB9 and the plasmid is absent in the accessory genome of iNTS strain D23580. This would suggest D23580 lacks both the *cib* and *imm* genes, and the implications of these observations have not been explored.

In this study, we confirmed that iNTS *S*. Typhimurium D23580 lacks colicin Ib activity in vitro. Surprisingly we observed the pathogen appeared to be resistant to the action of colicin Ib, even without the cognate immunity protein. Further investigation revealed that a non-typhoidal *S*. Typhimurium strain SL1344, engineered to lack the immunity gene, also exhibited resistance. This suggests that the immunity gene may be functionally redundant in these strains. Additional research showed that this resistance could be partially explained by variations in the colicin Ib receptor, CirA. However, it was evident that other factors also contribute to the resistance. Further analysis indicated that variations in the TonB transporter did not affect resistance under the tested conditions.

## Materials and Methods

### Bacterial strains and culture conditions

The bacterial strains and plasmids used in this work are listed in **Tables 1** and **2**. Bacteria were grown overnight at 37°c in liquid Lennox Broth (LB) with shaking at 180rpm or statically on LB agar unless otherwise stated. When necessary, this was supplemented with antibiotics (ampicillin 50µg/ml, kanamycin 50µg/ml and chloramphenicol 20µg/ml).

**Table 1:**
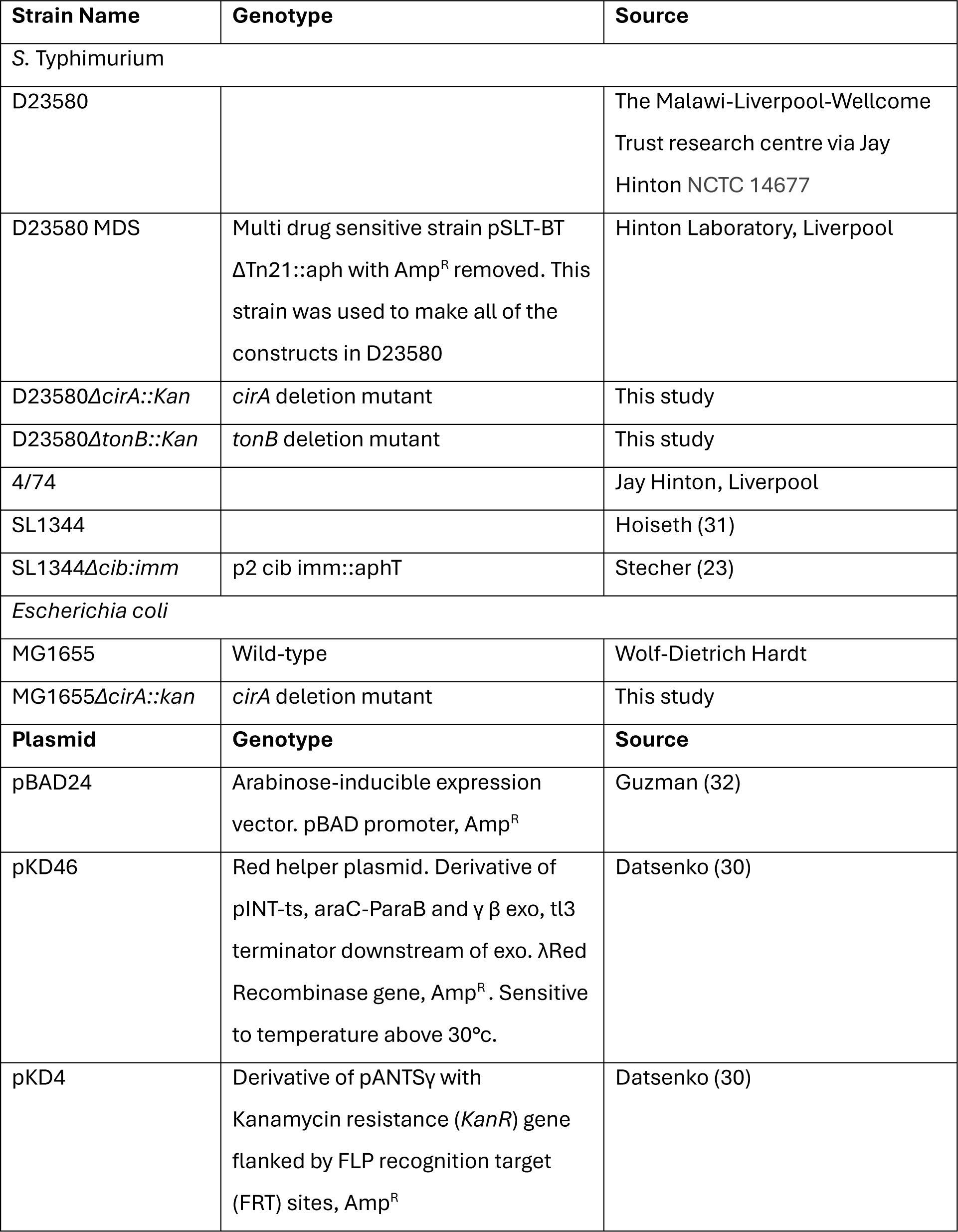
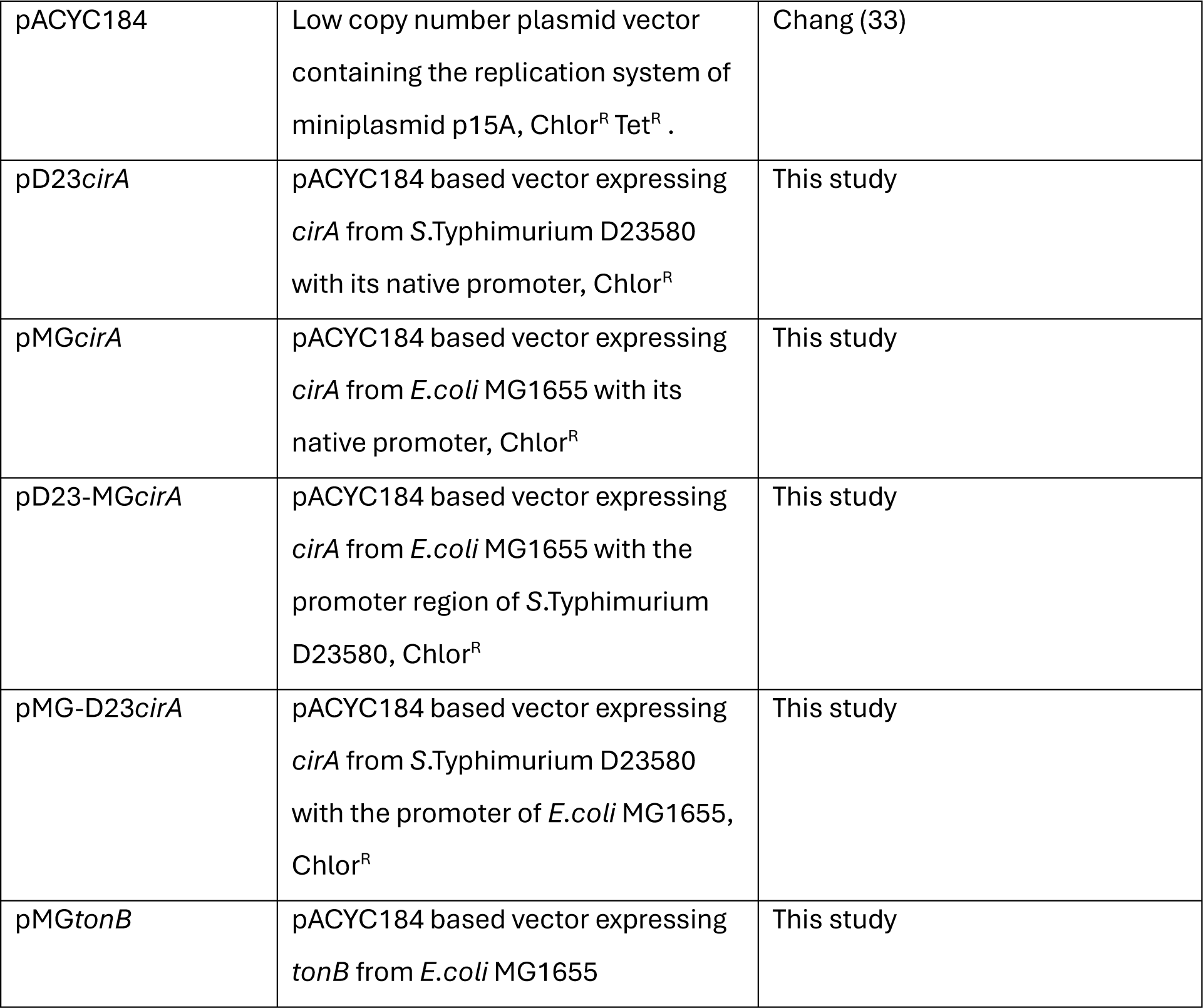
Bacterial strains and plasmids used in this study.

**Table 2:**
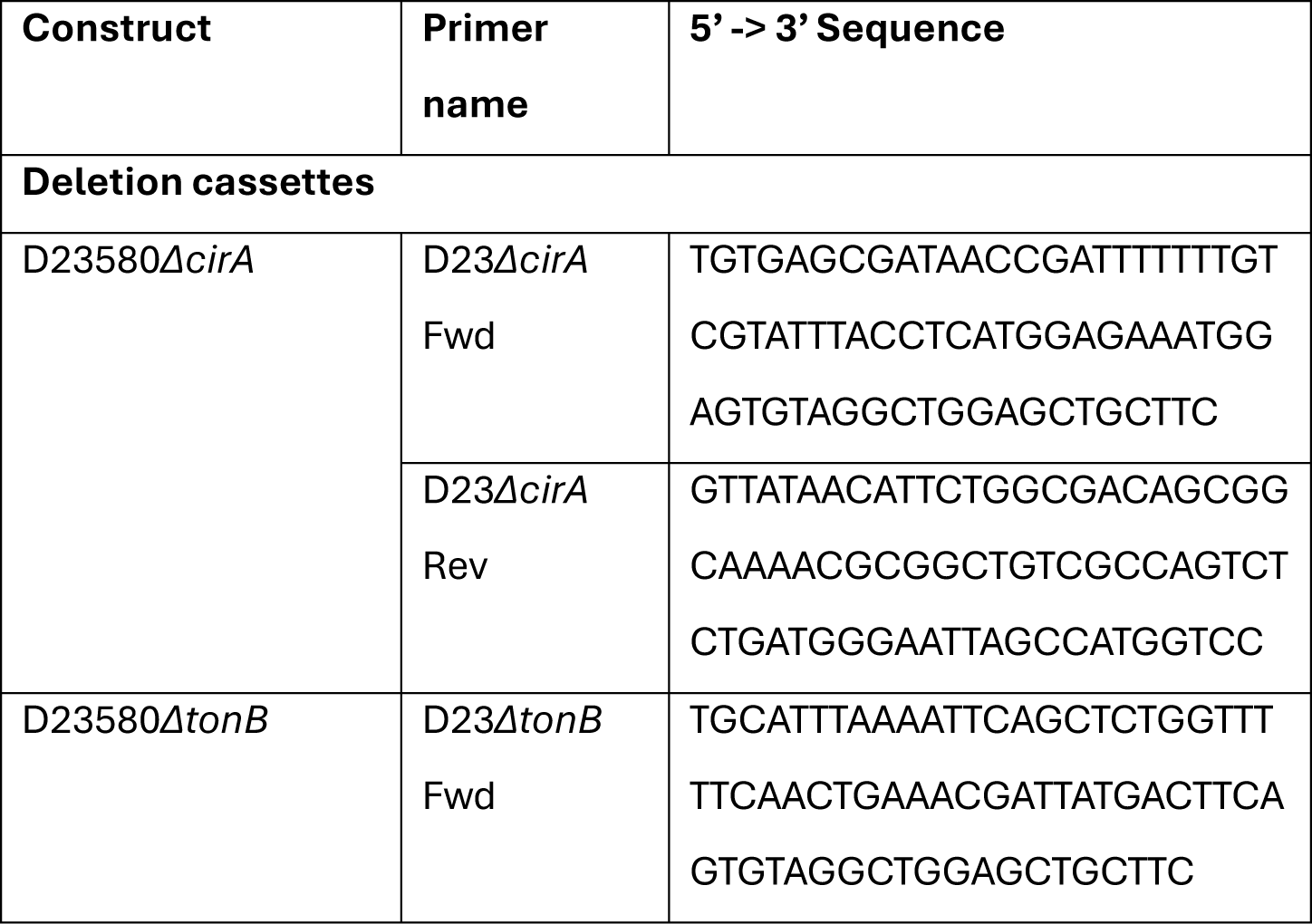
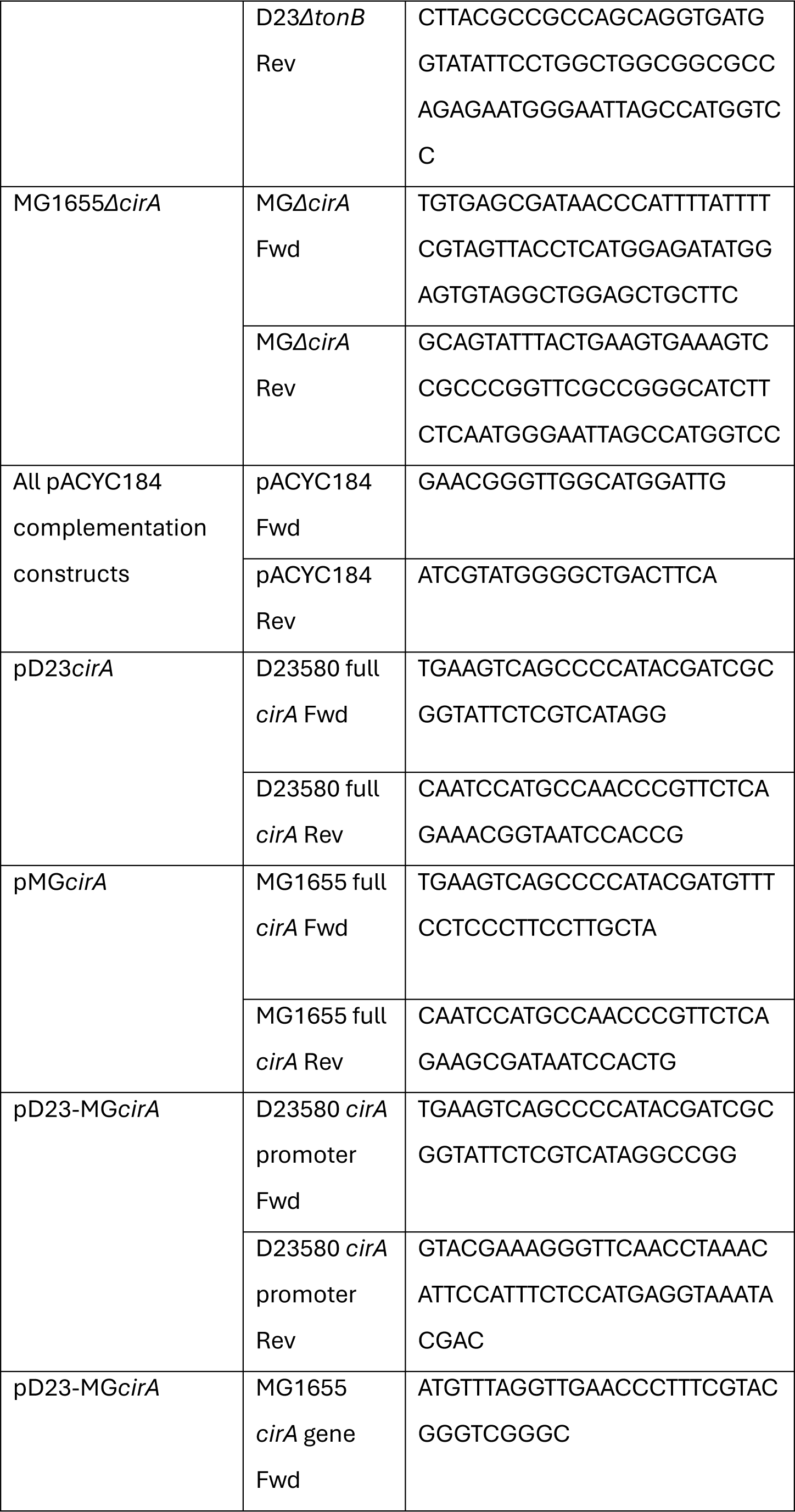
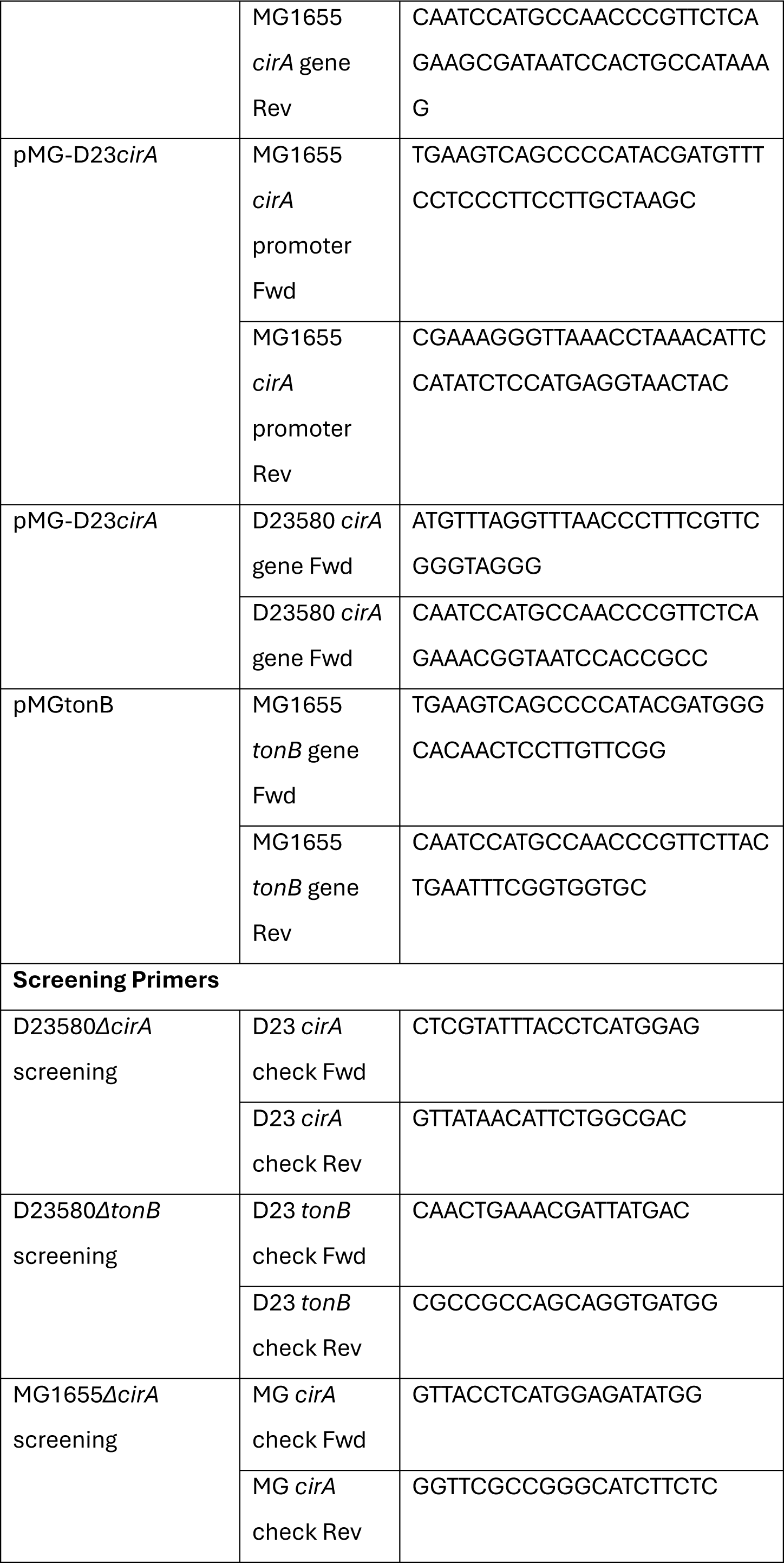
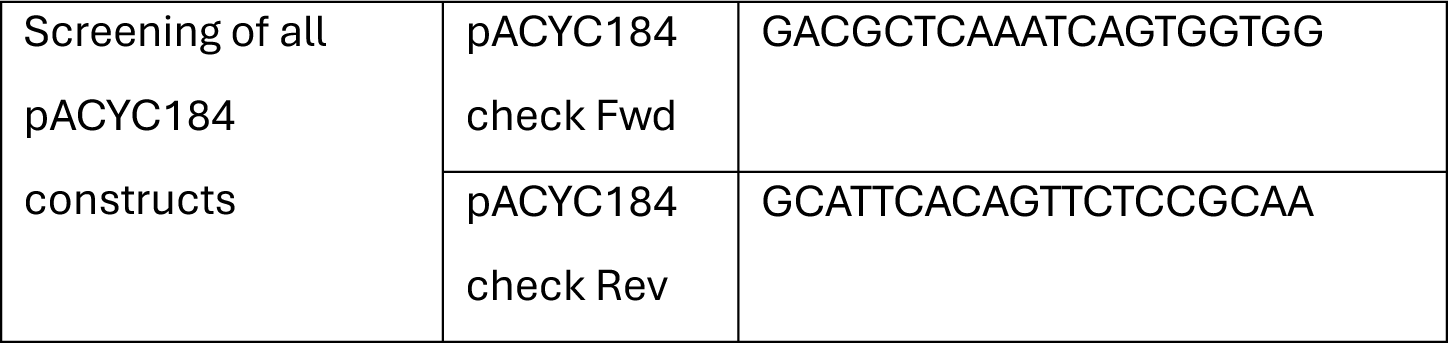
Primers used in this study.

### Construction of bacterial mutants and plasmids

All bacterial mutants and plasmids generated in this study are listed in **Table 1**. *E. coli* MG1655 and *S*. Typhimurium D23580 *ΔcirA* and *ΔtonB* were constructed using the lambda Red recombinase system described by Datsenko (30), using pKD4 as a template for the kanamycin-resistant cassette with flanking FRT-sites. The amplification primers used to generate these cassettes are shown in **Table 2**.

pACYC184 based complementation plasmids containing variants of the *cirA* gene were generated using Gibson assembly to combine DNA fragments PCR amplified using primers detailed in **Table 2**. Primers MG1655-CirA Fwd and Rev, and D23580-CirA Fwd and Rev were used to amplify *cirA* from the chromosomal DNA of MG1655 and D23580 along with their natural promoters, where flanking regions homologous to the insert site on pACYC184 were included on the primers. Promotor regions and genes were also amplified separately from both strains to generate plasmids pMG-D23 cirA and pD23-MG cirA.

Four microliters of Gibson assembly mix were transformed into chemically competent *E. coli* NEB5α (New England Biolabs) prior to transformation into the appropriate electrocompetent strains. Constructs were verified using colony PCR with screening primers detailed in **Table 2** followed by sequencing (Eurofins).

### Analysis of growth in co-culture

Co-cultures between *S.* Typhimurium and *E. coli* (1:1) utilised a 1:50 inoculation of LB, where 100 µM chelating agent DTPA (Diethylenetriamine pentaacetate) was added to induce colicin production, as described by Nedialkova (14). CFU/ml was determined following growth on selective agar (MacConkey agar).

### Analysis of growth in conditioned media

In order to assess bacterial growth over time, OD_600_ was analysed in a Tecan Sunrise plate reader. 96 well, clear plates contained 200 µl of media inoculated 1/50 with the desired bacteria. Growth was performed at 37 °c, with shaking, for 16 hours with measurements taken approximately every 5 minutes. Data analysis was performed using Graphpad Prism software.

As standard, conditioned media was produced from an overnight culture (∼16 hours). 100 µM chelating agent DTPA was added prior to overnight culture. Cells were then pelleted by centrifugation at 4 °c, 3,000 RPM, 10 minutes. The resulting supernatant was then enriched with 10x media to a final concentration of 1x before being filter sterilised through a 0.22 µm syringe filter. The conditioned media was then inoculated, and growth analysed as described above.

### Screening for the presence of *cib and imm* genes in *S. enterica* and *S.* Typhimurium

A total of 870 complete genomes of *S. enterica*, including 142 serovars, were downloaded from Enterobase (34). A minimum spanning tree was produced and visualised using GrapeTree v1.5.0 (35). The presence and absence of the *cib* and *imm* genes in the genomes were searched using BLASTN (36) with 80 % identity and 80% coverage.

A total of 1840 draft genomes and 6 complete genomes of *S.* Typhimurium were used to investigate the presence of *cib* and *imm* genes using BLASTN (80% identity and 80% coverage) (36). Using the core genome multi-locus sequence typing (cgMLST) scheme of *S.* Typhimurium (34, 37), allele profiles for the 1846 genomes were produced using chewBBACA v2.8.5 (38). Using the allele profile as input, a neighbor-joining tree was constructed using GrapeTree v1.5.0 (35). The tree was visualised using iTOL v6 (39).

## Results

### The presence of colicin Ib genes in *Salmonella* serovars

Although research, such as that by Nedialkova (14) has demonstrated the competitive advantage attributed to colicin Ib in select *S*. Typhimurium strains, it is unclear the proportion of *Salmonella* serovars that possess *cib* and *imm* and therefore benefit from the colicin.

Analysis of 870 complete *Salmonella* genomes of various serovars is shown in **Figure 2** with each serovar represented as a bubble on the minimum spanning tree. The presence of *cib* and *imm* is highlighted within each serovar in panel b. The majority of *Salmonella* serovars do not contain *cib* and *imm*, with the presence largely limited to serovars Typhimurium, Heidelberg, Anatum and Newport. Notably the *cib* and *imm*, were largely absent in systemically invasive serovars such as Typhi and Paratyphi, although it was also absent in gastrointestinal serovars such as Enteritidis and Dublin. Percentage breakdowns of serovars containing *cib* and *imm* are shown in **Table 3**.

**Figure 2:**
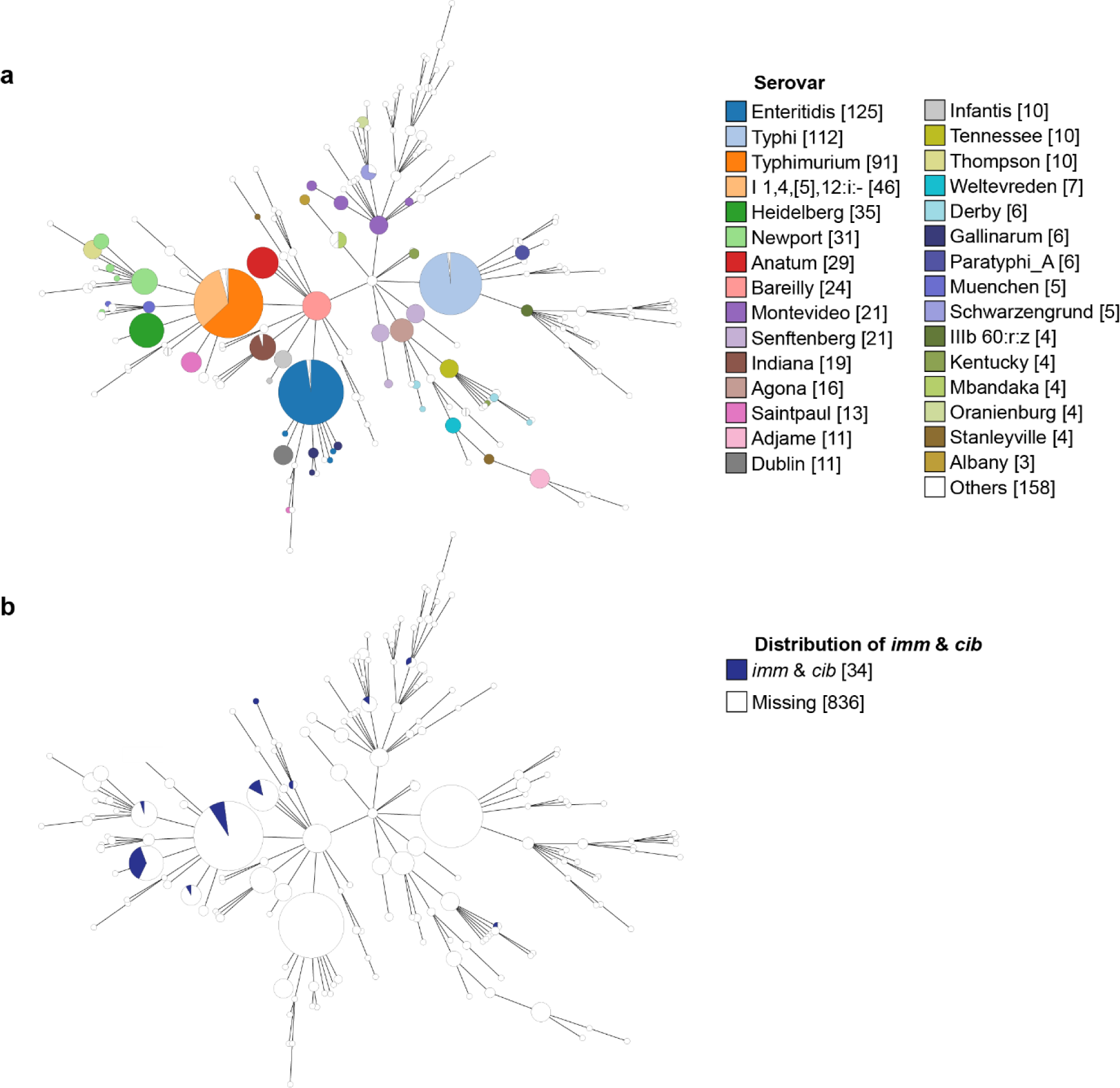
Presence of *imm* and *cib* in 870 *Salmonella* complete genomes. a. minimum spanning tree of genomes as visualized in GrapeTree, where different colours represent different serovars and the bubble size indicates the number of genomes. b. 34 genomes carry both *imm* and *cib*, denoted in blue.

**Table 3.** Distribution of *cib* and *imm* genes in different *Salmonella* serovars.

| Serovars | No. of strains |  | % *cib* & *imm* (+) |
| --- | --- | --- | --- |
|  | *cib* & *imm* (+) | Total |  |
| Total | 34 | 870 | 4% |
| Heidelberg | 13 | 35 | 37% |
| Anatum | 4 | 29 | 14% |
| Typhimurium | 10 | 91 | 11% |
| Saintpaul | 1 | 13 | 8% |
| Newport | 1 | 31 | 3% |
| Ohio | 1 | 2 | 1/2 |
| Paratyphi B var. Java | 1 | 2 | 1/2 |
| Rissen | 1 | 2 | 1/2 |
| Brandenburg | 1 | 3 | 1/3 |
| Schwarzengrund | 1 | 5 | 1/5 |

### The distribution of colicin Ib (*cib*) and its cognate immunity protein (*imm*) across the various strains of *S*. Typhimurium

As demonstrated in **Figure 2**, 11% of the Typhimurium genomes analysed carried the *cib* and *imm* genes. To better understand the distribution, a further breakdown within this serovar is shown in **Figure 3**. Analysis focused on 1840 strains of *S.* Typhimurium including multi-locus sequence types ST19 (associated with gastroenteritis) and ST313 lineages 1-3 (associated with invasive non-typhoidal Salmonellosis).

**Figure 3.**
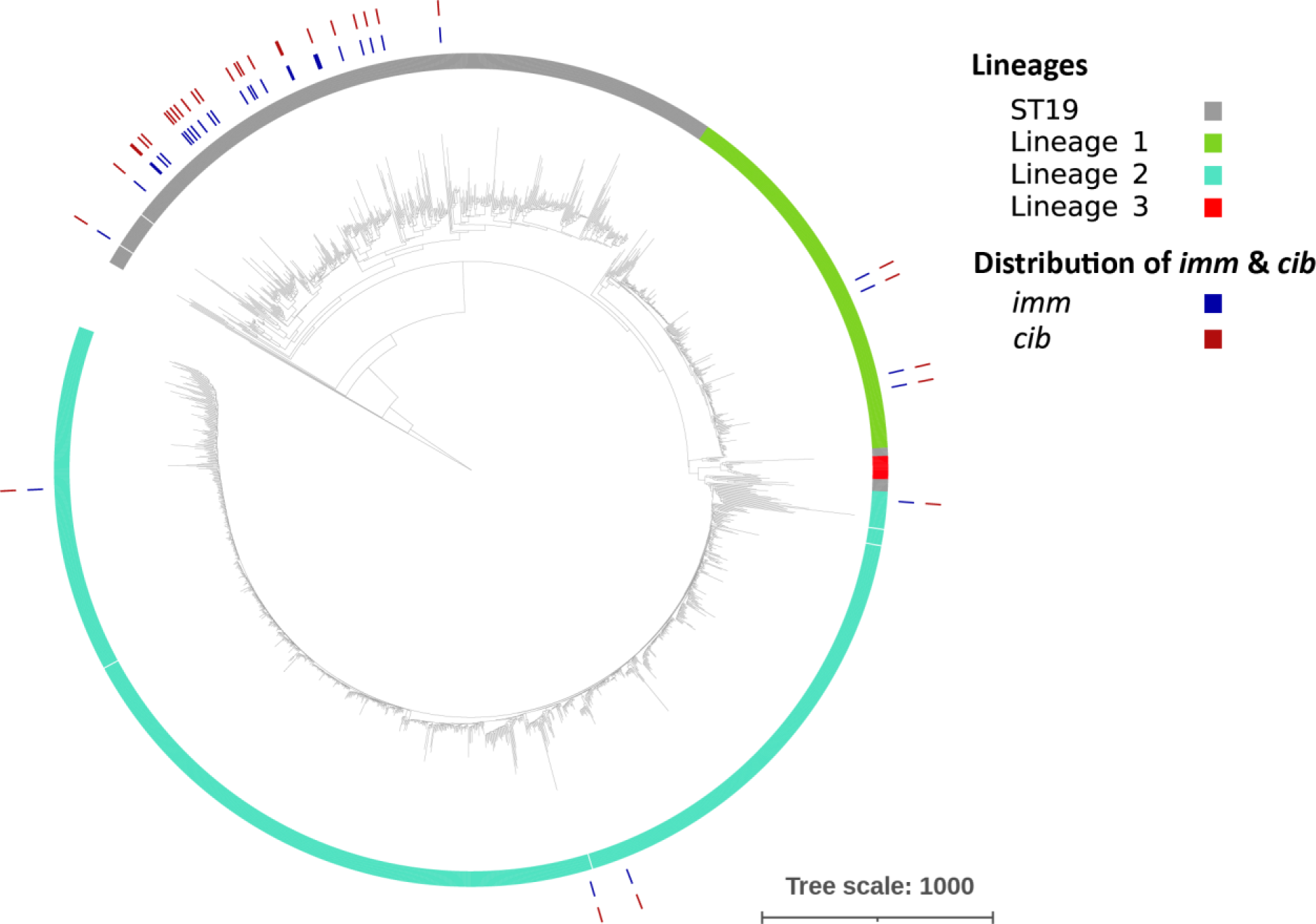
Presence of *imm* and *cib* in 1840 *S*. Typhimurium strains. The inner ring represents the population structure, with lineages 1 - 3 belonged to ST313, which are associated with iNTS. The ST19 strains are associated with gastroenteritis. The outer rings indicate the presence of *cib* and *imm*.

**Figure 4.**
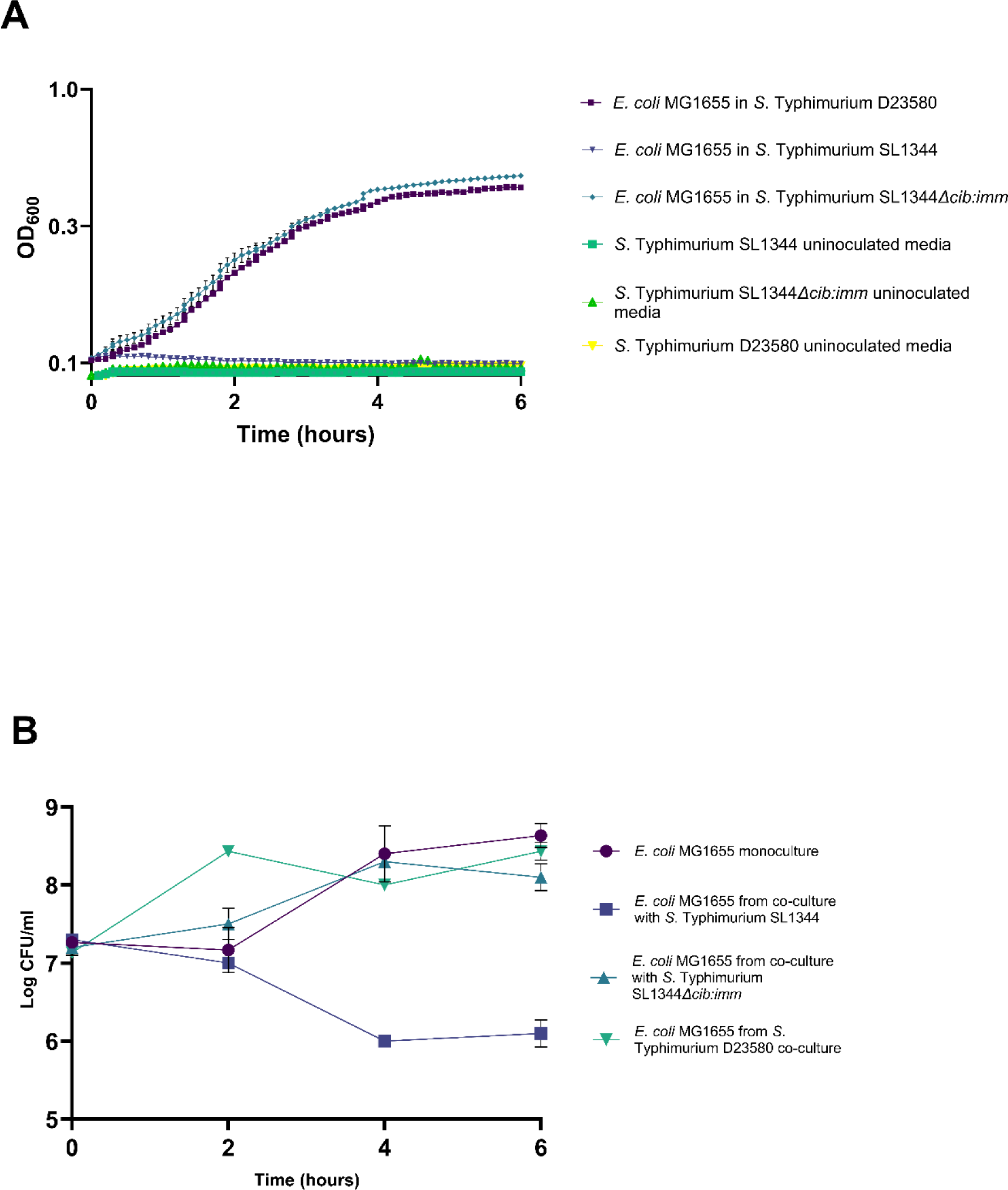
Growth of *E. coli* MG1655 in conditioned media from *S*. Typhimurium strains SL1344, SL1344Δ*cib:imm* and D23580. Growth kinetics of *E. coli* MG1655 over 6 hours in LB and conditioned media from *S*. Typhimurium strains grown in the presence of DTPA (panel A). Panel B shows the log CFU/ml of *E. coli* MG1655 in co-culture with *S*. Typhimurium strains compared to in monoculture. Data represent average values of triplicate measurements are shown +/-standard deviation where independent replicates were also performed (n=4). Generated using Graphpad Prism®.

It is clear that the colicin Ib associated genes, *cib* and *imm* are more commonly possessed by ST19 strains (4.9%) than those within ST313 (0.6%). Within ST313, the presence of *cib* and *imm* decreased with each lineage (1.4% of lineage 1 strains, 0.4% of lineage 2 and 0% of lineage 3). Interestingly, two of the genomes tested from ST19 contained the *imm* gene but not the *cib*.

Figures 2 and **3** suggest that the colicin Ib immunity protein is absent from a significant proportion of *Salmonella*, but the implications of this in the colicin rich gastrointestinal environment has not been investigated.

### iNTS strain *S*. Typhimurium D23580 does not produce Colicin Ib

Genomic analysis of *S*. Typhimurium D23580 (ST313, lineage 2 iNTS strain) did not identify *cib* or *imm* genes within the genome of the strain. The effects of colicin Ib on *imm* deficient strains such as D23580 has not previously been explored.

**Table 4.** Distribution of *cib* and *imm* genes in *S*. Typhimurium.

| Lineages | No. of strains |  |  | % of *cib* & *imm* (+) |
| --- | --- | --- | --- | --- |
|  | *cib* & *imm* (+) | *imm* only | Total |  |
| ST19 | 25 | 2 | 514 | 4.9% |
| ST313 - Lineage 1 | 4 |  | 276 | 1.4% |
| ST313 - Lineage 2 | 4 |  | 1039 | 0.4% |
| ST313 - Lineage 3 |  |  | 17 | 0.0% |
| Total | 33 | 2 | 1846 | 1.8% |

*E. coli* MG1655 is known to be highly sensitive to colicin Ib, and therefore was selected as a control by which colicin activity could be determined. The non-typhoidal *S.* Typhimurium strain 4/74 harbours both the *cib* and *imm* genes. Chelating agent DTPA (Diethylenetriamine pentaacetate) was added to induce colicin production, as described by (14). In vitro growth analysis of *E. coli* MG1655 in spent media from a panel of *S.* Typhimurium strains, including *S*. Typhimurium D23580 showed no evidence of colicin production in the iNTS strain, as denoted by growth comparable to that in control media. In contrast, colicin dependent growth repression, as identified in *S.* Typhimurium strain 4/74 WT media compared to 4/74 *Δcib:imm* mutants, was identified in the colicin producing NTS strain.

These observations were also consistent with the respective co-cultures, where reduced MG1655 was seen in co-culture with colicin producing NTS bacteria but not with *Δcib:imm* mutants or iNTS.

### *S*. Typhimurium is resistant to colicin Ib in the absence of the immunity gene, *imm*

Colicin Ib bacteriocidal activity is associated with *E. coli* as the target organism, with little research into the sensitivity of *S*. Typhimurium in the absence of the immunity gene. Research by Schneider (40) showed (supplementary data, figure 1) that from a panel of 35 *S. enterica* serotypes, including two Typhimurium strains, 90% of the strains tested showed sensitivity to colicin Ib. Although the immunity gene is often referred to as protecting colicin Ib-producing strains of *Salmonella*, the importance of this gene has not been fully investigated.

In order to investigate the importance of the immunity gene *imm* in *S*. Typhimurium, invasive non-typhoidal strain D23580 was utilised (lacks the colicin Ib and immunity proteins), as well as a Δ*cib:imm* mutant from colicin producing strain *S*. Typhimurium SL1344, which also lacks the colicin production and immunity genes. Growth of these strains, in colicin rich spent media of producing strains (*S*. Typhimurium 4/74 and SL1344) would determine if the immunity gene is vital in the defence of Typhimurium strains against colicin Ib.

Both *imm* deficient strains appeared to grow comparably in colicin rich conditioned media to growth in LB only, showing optimal growth under the conditions tested (figure 5). This suggests that these strains are tolerant to the effects of colicin Ib, despite lacking the immunity gene associated with protection.

**Figure 5.**
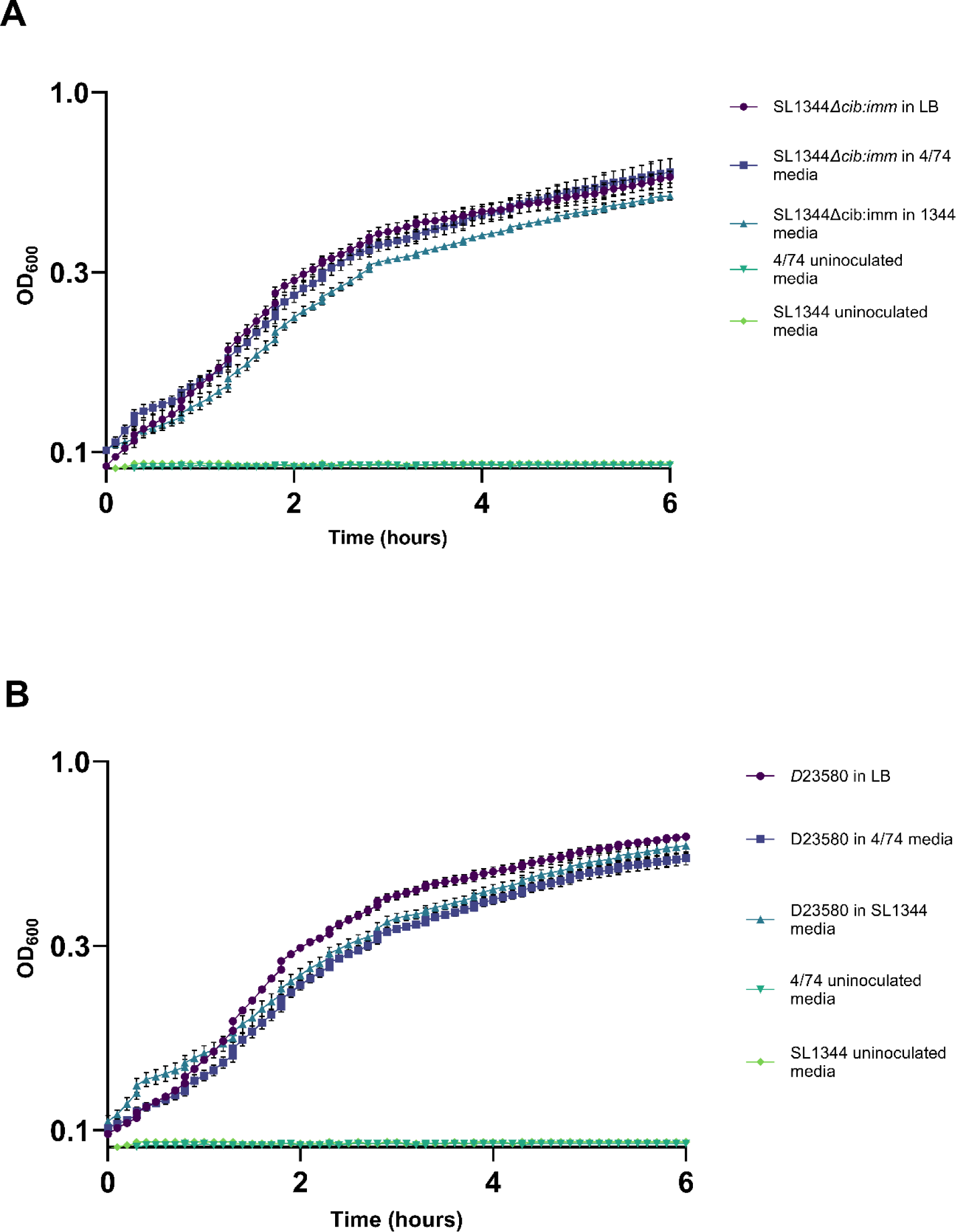
Growth analysis of *S*. Typhimurium D23580 and SL1344 *Δcib:imm* in colicin rich media. 6 hour growth kinetics of SL1344 *Δcib:imm* (panel A) and D23580 (panel B) in LB and conditioned media of colicin producing 4/74 and SL1344 strains grown in DTPA. Data represents the average of triplicate measurements +/-standard deviation where independent replicates n=3. Generated using Graphpad Prism®.

Tolerance to this colicin in target strains lacking immunity has been shown previously in *E. coli*, where mutations in the colicin receptor CirA are associated with resistance (14, 41).

It was hypothesised that the seeming intrinsic resistance of the *S*. Typhimurium strains tested may be due to variations in colicin receptor CirA from that of sensitive *E. coli* MG1655. Comparison of the amino acid sequences for the colicin receptor from both strains showed 88% similarity. The variation between these strains is shown in figure 6, where alternative amino acids of *S*. Typhimurium D23580 are mapped onto the structure of *E. coli* CirA.

**Figure 6.**
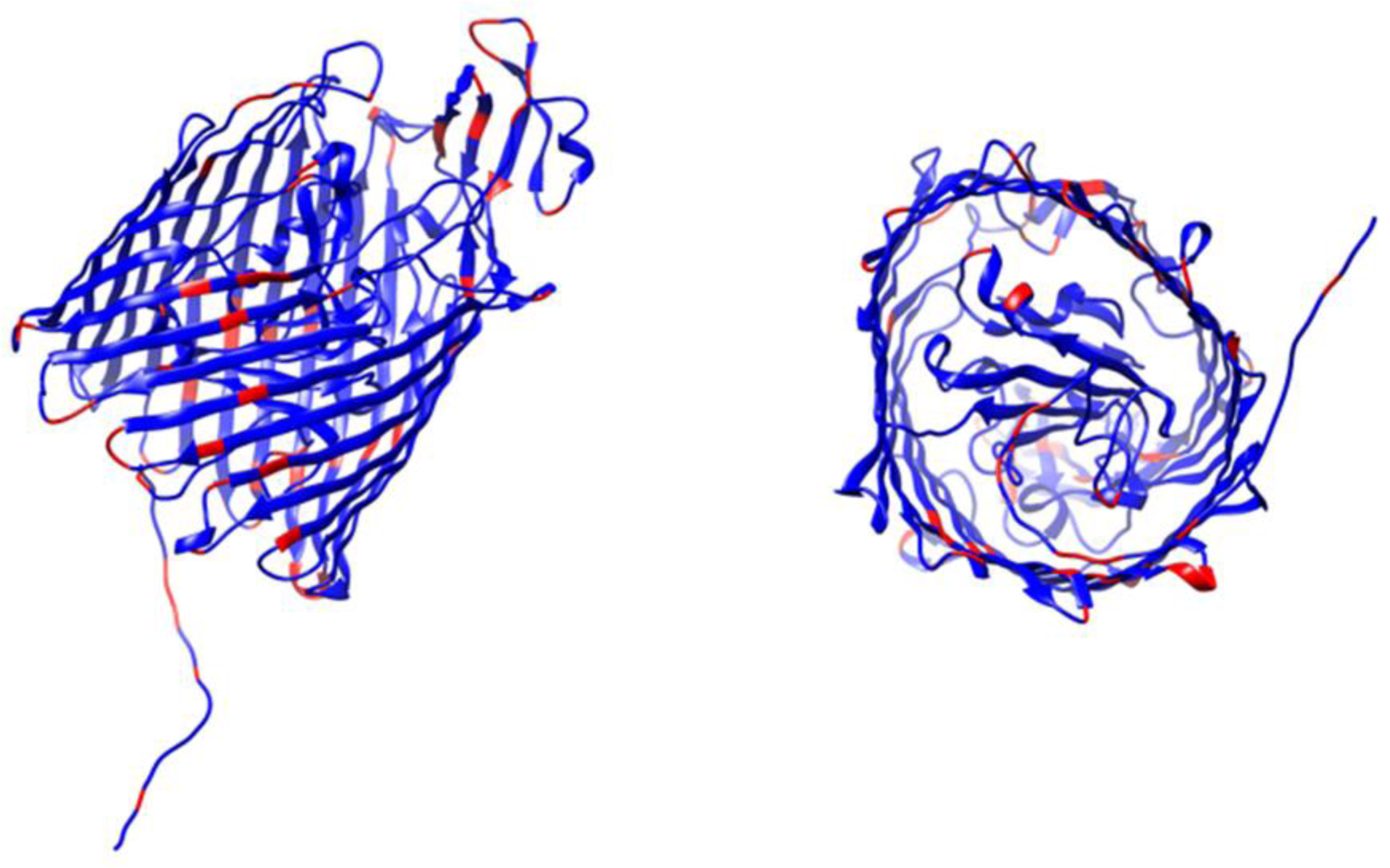
The CirA protein of *E. coli* is shown in blue, where amino acids substitutions found in *S*. Typhimurium D23580 are highlighted in red. The protein is shown in multiple orientations, where this figure was constructed from the published crystal structure PDB 2HDF with annotation in Chimera®

In order to determine if the variation in CirA contributed to the resistance to colicin Ib in *S*. Typhimurium, specifically iNTS *S*. Typhimurium D23580, deletion mutants were complemented with the converse *cirA* using plasmids pMG*cirA* and pD23*cirA* shown in **Table 1**. The effects of this substitution on colicin sensitivity could then be assessed by observing the growth of the complemented mutants in colicin rich media.

When the colicin receptor *cirA* of previously colicin resistant *S*. Typhimurium D23580 is replaced by *cirA* of sensitive *E. coli* MG1655, a shift in the growth in colicin rich (*S*. Typhimurim SL1344) media is observed (Figure 7A). Compared to that of WT D23580 in the same media, the complemented mutant shows reduced growth, suggestive of a degree of colicin sensitivity being introduced. This altered growth was determined to be colicin dependent by the growth in SL1344*Δcib:imm* conditioned media (lacking colicin Ib), where no reduction in growth was evident. The growth of this complemented mutant in LB is comparable to that of the WT strain, suggesting that the mutation of *cirA* and burden of the complementation plasmid, is not affecting the growth fitness of the strain.

**Figure 7:**
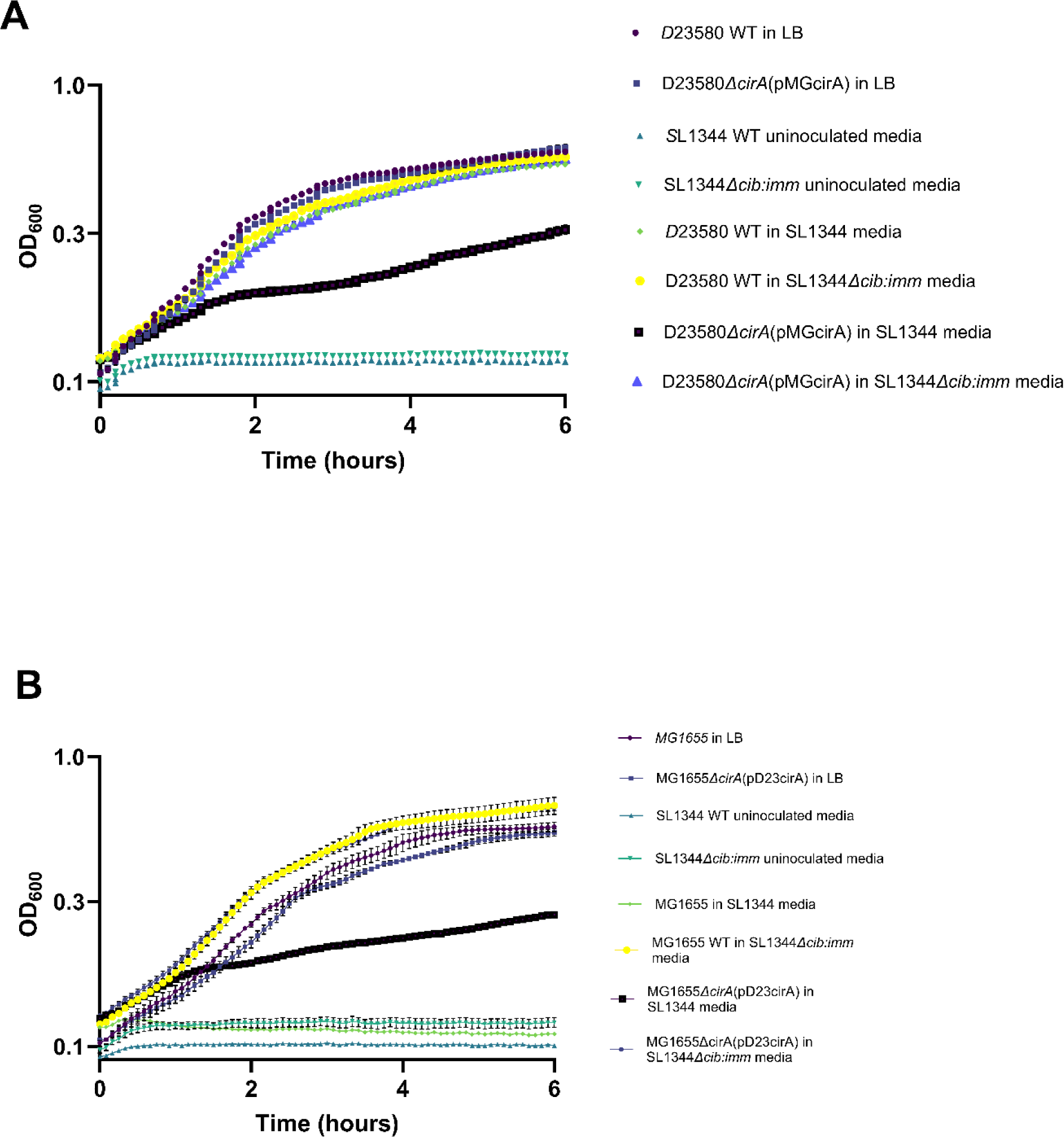
Growth of complemented colicin mutants *S*. Typhimurium D23580*ΔcirA*(pMGcirA) and *E. coli* MG1655*ΔcirA*(pD23cirA) in the presence and absence of colicin Ib. 6 hour growth kinetics of *S*. Typhimurium D23580*ΔcirA*(pMGcirA) and *E. coli* MG1655*ΔcirA*(pD23cirA) in panels A and B respectively. Growth LB and in conditioned media from *S*. Typhimurium SL1344 and *S*. Typhimurium SL1344*Δcib:imm* grown with DTPA is shown. Data represents the average of triplicate measurements +/-standard deviation where independent replicate n=3. Generated using Graphpad Prism®.

To further characterise the role of CirA in the resistance of *S*. Typhimurium D23580 to colicin Ib, the role of the receptor regulation was investigated. In order to separate the roles of both the regulation of the gene, and the gene itself, hybrid plasmids with “swapped promoters” were constructed in the relevant *ΔcirA* strains. This included the *cirA* gene from resistant strain *S*. Typhimurium D23580 under the control of the *cirA* promoter region from *E. coli* MG1655 and vice versa.

Compared to WT *S*. Typhimurium D23580, the introduction of the *E. coli* MG1655 *cirA* gene under the control of the D23580 promoter (pD23-MG *cirA*) increased sensitivity to the colicin, showing reduced growth (Figure 8A). In contrast, expression of the *S*. Typhimurium D23580 *cirA* gene under the control of the *E. coli* MG1655 promoter (pMG-D23 *cirA*) did not appear to alter the growth of D23580*ΔcirA* compared to that of WT (Figure 8B). This suggests that the effects are dependent on the CirA structure rather than any altered regulation, consistent with the hypothesis that the resistant phenotype is associated with amino acid variation shown in *S.* Typhimurium strains.

**Figure 8:**
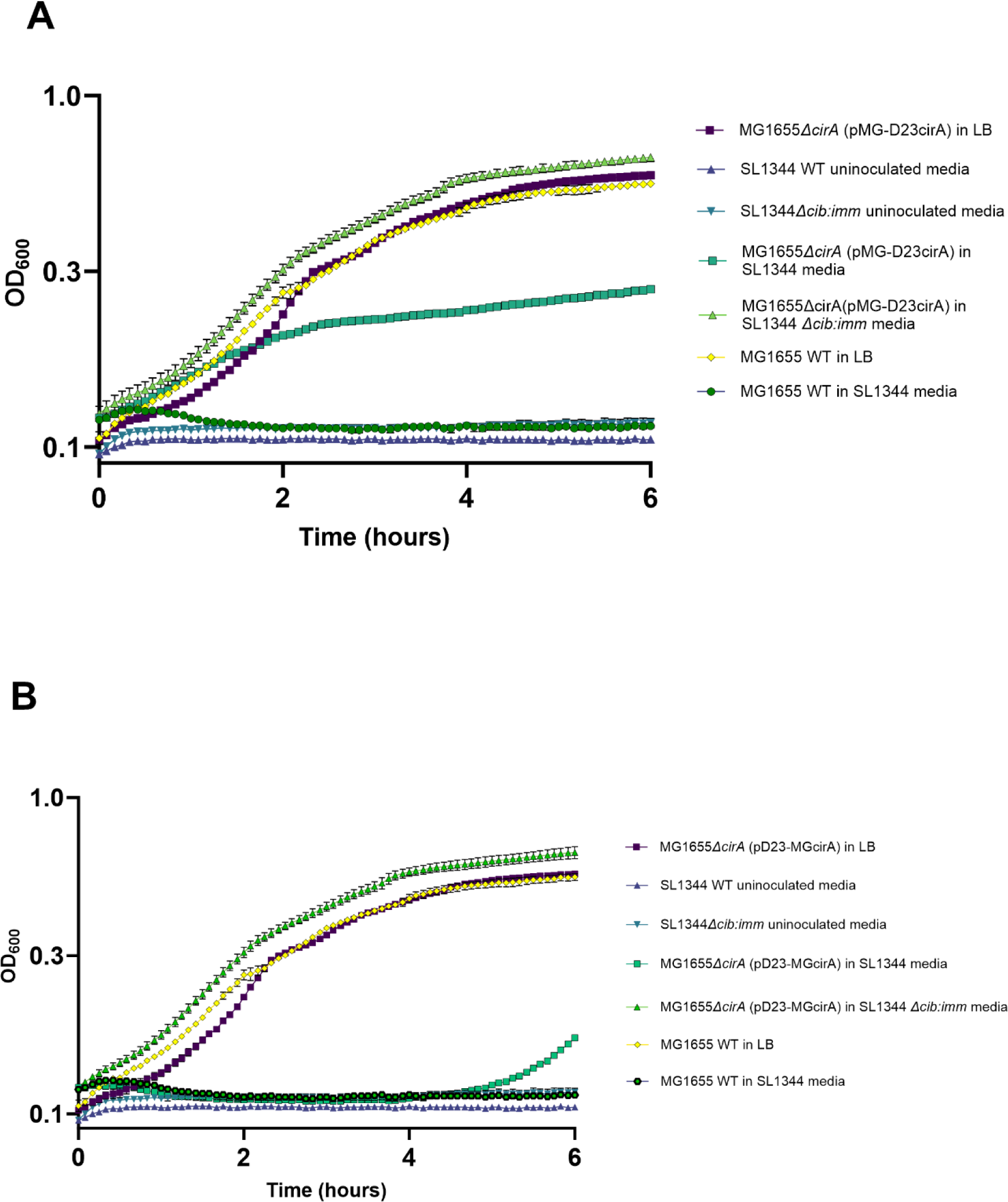
Growth analysis of *E. coli* MG1655*ΔcirA* complemented with pMG-D23cirA and pD23-MGcirA in the presence and absence of colicin Ib. 6 hour growth kinetics of *E. coli* MG1655*ΔcirA* pMG-D23cirA and pD23-MGcirA in panels A and B. Growth LB and in enriched, spent media from *S*. Typhimurium SL1344 WT and SL1344*Δcib:imm* grown with DTPA is shown. Averages of triplicate measurements +/-standard deviation where independent replicate n=3. Generated using Graphpad Prism®.

Consistently, introducing the *cirA* gene of *S*. Typhimurium D23580 under the control of the *E. coli* MG1655 promoter (pMG-D23), where a reduction in sensitivity was shown in comparison to WT *E. coli* MG1655 (**Figure 9A**. In contrast to the findings in *S*. Typhimurium D23580, introduction of pD23-MG to *E. coli* MG1655*ΔcirA* introduced altered growth compared to that of WT bacteria, where regulation by the Typhimurium promoter appeared to reduce the effects of colicin Ib, despite the presence of *cirA* from the sensitive strain (Figure 9B).

**Figure 9:**
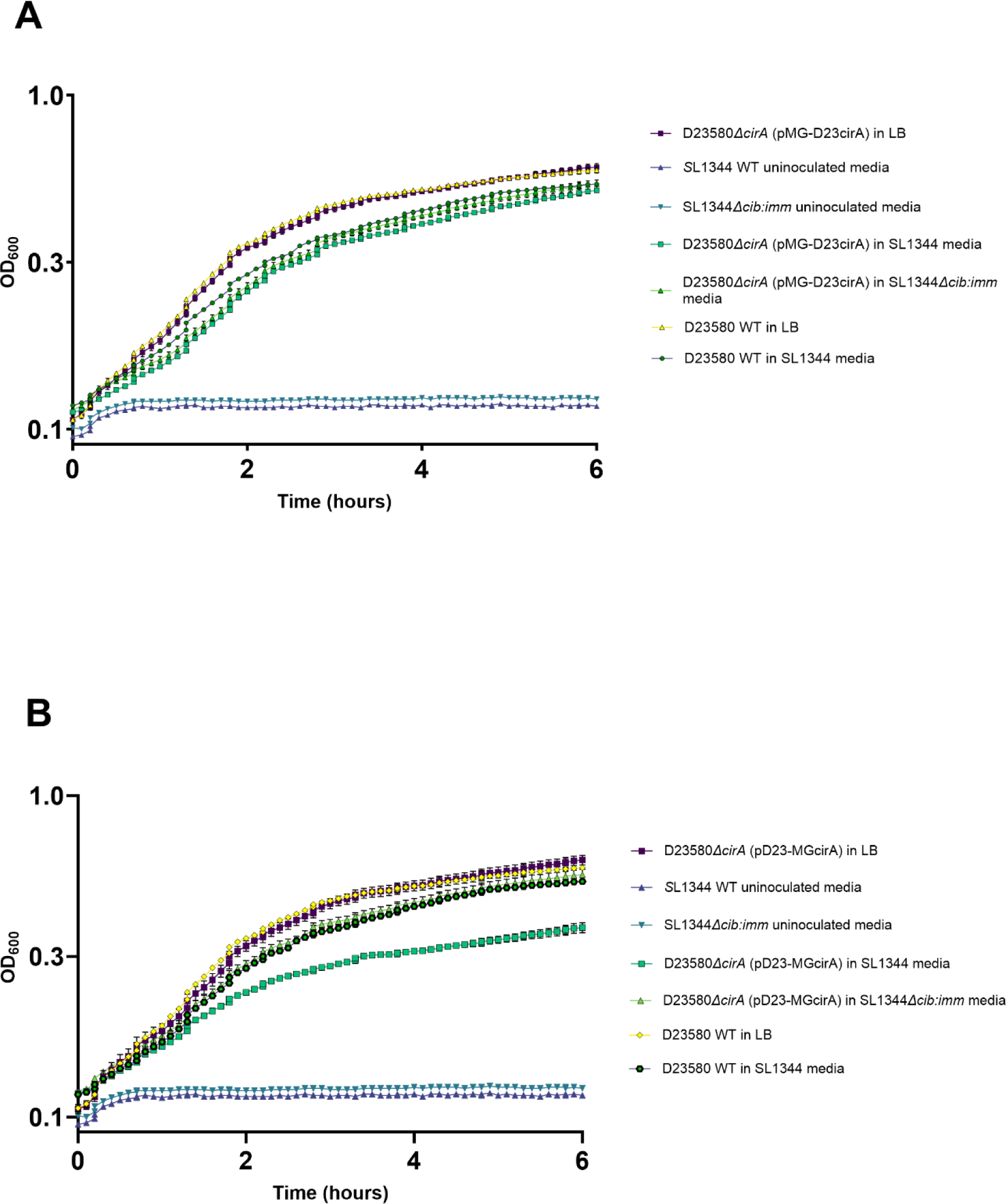
Growth analysis of *S*. Typhimurium D23580*ΔcirA* complemented with pMG-D23cirA and pD23-MGcirA in the presence and absence of colicin Ib. 6 hour growth kinetics of *S.* Typhimurium D23580*ΔcirA* pMG-D23cirA (panel A) and pD23-MGcirA (panel B). Growth in LB and conditioned media from *S*. Typhimurium SL1344 WT and SL1344*Δcib:imm* grown with DTPA is shown. Averages of triplicate measurements +/-standard deviation where independent replicate n=3. Generated using Graphpad Prism®.

Although it is clear that colicin receptor, CirA, contributes to the resistance of iNTS strain *S*. Typhimurium D23580 to colicin Ib, the partial introduction of resistance in *E. coli* and sensitivity in iNTS respectively suggests that multiple factors are at play.

Following interactions with CirA, colicins Ia and Ib are transported to the inner membrane in a TonB dependent manor. Therefore, mutations in TonB are also associated with resistance to these colicins (25). It was hypothesised that the amino acid variation seen between these strains may contribute to the colicin resistance seen in iNTS.

As with *cirA*, Δ*tonB* deletion mutants were complemented with alternative *tonB* (*S*. Typhimurium with *E. coli tonB* and vice versa) were grown in colicin Ib rich media to assess the impact of the substitution on colicin sensitivity.

Unlike *cirA*, the introduction of *E. coli* MG1655 *tonB* into *S*. Typhimurium D23580 did not introduce any sensitivity to the colicin under the conditions tested. Additionally, the presence of D23580 *tonB* in MG1655 did not provide any resistance to colicin Ib as was seen when *cirA* was substituted.

In combination, this suggests that the genetic variation of TonB between the colicin resistant *S*. Typhimurium and sensitive *E. coli* is not a contributing factor to the resistance of *Salmonella* under the conditions tested.

**Figure 10:**
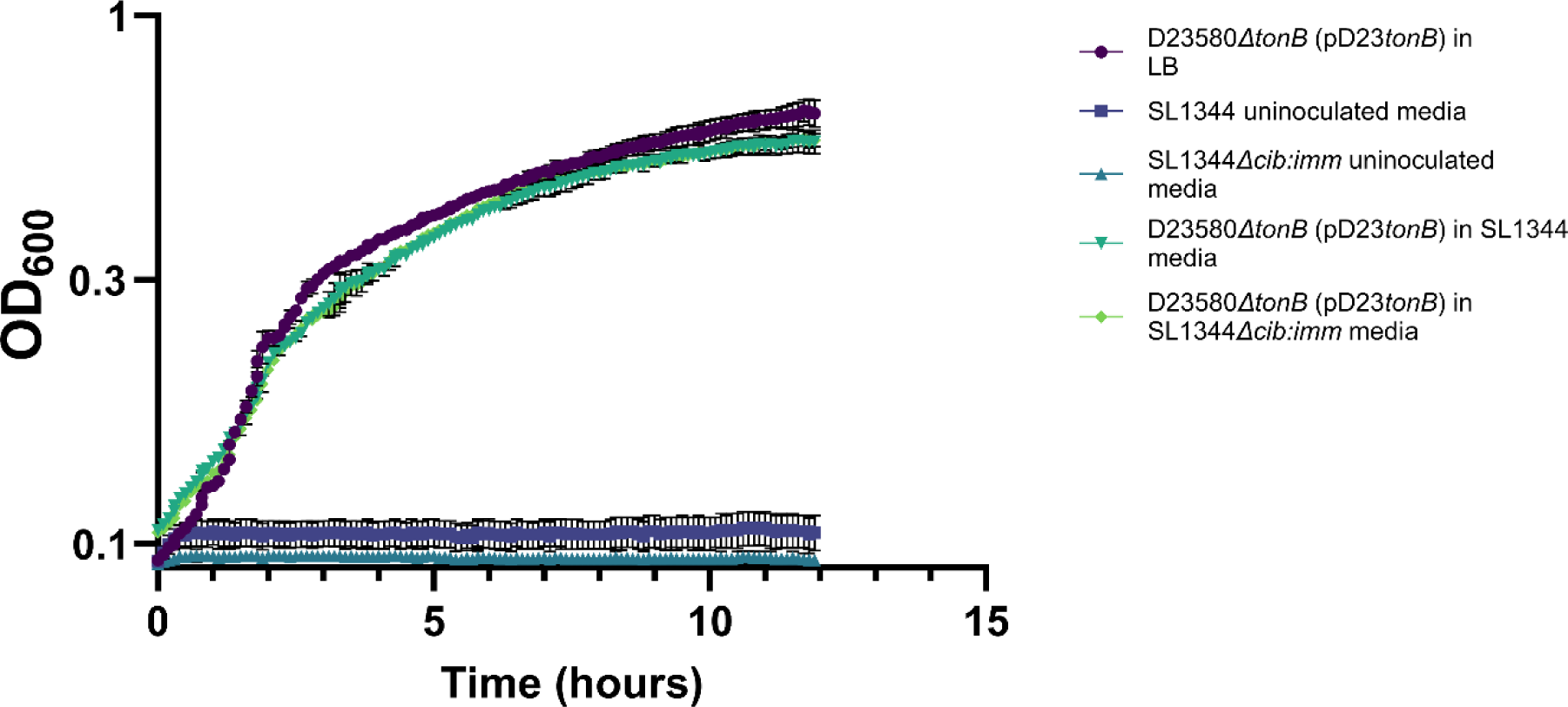
Growth analysis at of D23580*ΔtonB* complemented with pMG*tonB* in the presence and absence of colicin Ib. 6 hour growth kinetics of D23580Δ*tonB* complemented with *tonB* from *E. coli* MG1655. Growth in LB and in conditioned media from *S*. Typhimurium SL1344 WT and SL1344*Δcib:imm* grown with DTPA. The average of triplicate measurements +/-standard deviation is shown, where independent replicate n=3. Generated using Graphpad Prism®.

## Discussion

The role of colicins in colonisation resistance the prevention of colonisation of invading pathogens by the resident microbiota, is a fascinating and expanding field of research. The studies have focussed on the effects of these potent bacteriocins on commensal *E. coli* and the competitive edge they confer. It is thought that the presence of the pColIb plasmid in non-typhoidal *Salmonella* strains 4/74 and SL1344 may provide an advantage against competing *Enterobacteriaceae* (42).

It has been well established that some strains of *S*. Typhimurium are able to produce colicin Ib due to the carriage of *cib* gene. The immunity gene, *imm*, has been proven to provide protection to colicin Ib producing strains (16, 43, 44).

iNTS strains have recently emerged through a process of reductive evolution, with genes important for intestinal colonisation acquiring mutations and becoming redundant pseudogenes (45, 46). It is interesting that the majority of iNTS strains, including D23580 lacks both the ColIb and immunity genes (*cib* and *imm*). This could be attributed to the fact that these iNTS strains cause systemic disease, rarely with gastrointestinal symptoms, compared to NTS gastroenteritis causing strains (18). Thus possessing these genes does not provide a competitive advantage for iNTS strains (47, 48). This is consistent with the lack of the colIb and immunity genes in the genomes tested from invasive serovars Typhi and Paratyphi genomes (with the exception of a single Paratyphi B var. Java genome), as well as the rare possession of the genes in iNTS associated ST313 compared to that of NTS associated ST19. The absence of the genes in gastrointestinal associated serovars such as Enteritidis shows that not all NTS strains possess this competitive advantage.

The effects of this colicin on *S.* Typhimurium lacking this immunity gene such as iNTS strain D23580 are unclear, with the implications of colicin sensitivity in the colicin rich environment of the intestine unexplored.

As it lacks the plasmid encoding both the colicin (*cib*) and immunity genes (*imm*), that invasive non-typhoidal *S*. Typhimurium D23580, unlike its non-invasive counterpart 4/74, does not produce colicin Ib. The invasive nature of this pathogen may reduce the benefits of producing colicin Ib, as interactions with the microbiota are relatively short lived, in comparison to non-invasive strains such as *S*. Typhimurium 4/74. The lack of colicin production was confirmed with growth of sensitive *E. coli*.

Despite lacking the colicin Ib immunity gene, iNTS was shown to be resistant to colicin Ib during aerobic growth in conditioned media derived from a colicin Ib producing strain. Additionally, NTS with a deletion mutation of the *imm* gene was also shown to be colicin resistant, suggesting an intrinsic resistance in the *S.* Typhimurium strains tested, which has not been previously described.

TonB dependent colicin receptor CirA is a key component in colicin activity, where mutations in *cirA* have long since been associated with colicin resistance (25). Through the construction of a series of mutations and complementation assays, this research showed that amino acid sequence variations between CirA of *E. coli* and *S*. Typhimurium contributed towards the colicin resistance. Interestingly, it was evident that CirA was not solely responsible for this resistance, where substitution of the *cirA* gene introduced partial sensitivity and resistance to the colicin in *S*. Typhimurium and *E. coli* respectively. Additionally, substitution of TonB did not appear to alter colicin sensitivity. This suggests that an unknown factor, independent of CirA and TonB, is also involved in the intrinsic resistance to colicin Ib demonstrated in iNTS and NTS strains of *S*. Typhimurium in the absence of the colicin immunity gene, *imm*.

The biological implications in this resistance remains unclear, where it is possible that the competitive advantages of producing colicin Ib are negligible in invasive *Salmonella* such as iNTS, Typhi and Paratyphi compared to NTS strains with more prolonged interactions with the gut microbiota. Where the colicin resistance of iNTS *S.* Typhimurium may allow the survival of those colicin non-producing strains in the transient interactions within the colicin rich gastrointestinal environment.

Interestingly, this hypothesis may be corroborated by recent genomic comparisons between lineage two ST313 strains, *S*. Typhimurium D23580 (L2.0) and *S*. Typhimurium D37712, a member of the novel sublineage (L2.2) shown to be replacing lineage 2.0 organisms (49).

In this study, the accessory genomes of the representative strains were compared, with L2.2 strain D37712 possessing the pCol1B9 plasmid which usually encodes colicin Ib and its cognate immunity protein. However, in this iNTS strain, the colicin toxin-antitoxin system is absent from the plasmid. This may suggest that, as hypothesised, that the production of the colicin is less advantageous to these strains compared to NTS counterparts, with the intrinsic resistance to the colicin shown in *S.* Typhimurium strains in this study also making the immunity protein redundant.

These findings highlight a complex interplay in bacterial resistance to colicin Ib in the absence of the cognate immunity protein Imm, underscoring the intricacies of microbial interactions and colonisation resistance within the gastrointestinal environment.

## Author Contributions

The study was conceptualized by BG, JH, and CMAK. The methodology was designed by BG, JH, and CMAK. The experimental investigations were conducted by BG. LL conducted the genomic bioinformatic analysis, with contributions from YL. Data curation and analysis was performed by BG, JCC, BQ, AA. The manuscript was written by BG and reviewed and edited by LL, JCC, BQ, JH, and CMAK. The study was supervised by JH and CMAK. All authors reviewed the manuscript and approved the final draft.

## References

1. Libertucci J, Young VB. The role of the microbiota in infectious diseases. Nature Microbiology. 2019;4(1):35–45.

2. Sana TG, Flaugnatti N, Lugo KA, Lam LH, Jacobson A, Baylot V, et al. Salmonella Typhimurium utilizes a T6SS-mediated antibacterial weapon to establish in the host gut. Proceedings of the National Academy of Sciences. 2016;113(34):044–51.

3. Hockenberry AM, Micali G, Takács G, Weng J, Hardt W-D, Ackermann M. Microbiota-derived metabolites inhibit Salmonella virulent subpopulation development by acting on single-cell behaviors. Proceedings of the National Academy of Sciences. 2021;118(31):e2103027118.

4. Palmer LD, Skaar EP. Cuts Both Ways: Proteases Modulate Virulence of Enterohemorrhagic Escherichia coli. Mbio. 2019;10(1):10.1128/mbio.00115-19.

5. Hegarty JW, Guinane CM, Ross RP, Hill C, Cotter PD. Bacteriocin production: a relatively unharnessed probiotic trait? F1000Research. 2016;5.

6. Sibinelli-Sousa S, de Araújo-Silva AL, Hespanhol JT, Bayer-Santos E. Revisiting the steps of Salmonella gut infection with a focus on antagonistic interbacterial interactions. The FEBS Journal. 2022;289(14):4192–211.

7. Ducarmon Q, Zwittink R, Hornung B, Van Schaik W, Young V, Kuijper E. Gut microbiota and colonization resistance against bacterial enteric infection. Microbiology and Molecular Biology Reviews. 2019;83(3):10.1128/mmbr.00007-19.

8. Sassone-Corsi M, Nuccio S-P, Liu H, Hernandez D, Vu CT, Takahashi AA, et al. Microcins mediate competition among Enterobacteriaceae in the inflamed gut. Nature. 2016;540(7632):280-3.

9. Gradisteanu Pircalabioru G, Popa LI, Marutescu L, Gheorghe I, Popa M, Czobor Barbu I, et al. Bacteriocins in the era of antibiotic resistance: Rising to the challenge. Pharmaceutics. 2021;13(2):196.

10. Nawrocki EM, Hutchins LE, Eaton KA, Dudley EG. Mcc 1229, an Stx2a-amplifying microcin, is produced in vivo and requires CirA for activity. Infection and Immunity. 2022;90(2):e00587–21.

11. Mathur H, Field D, Rea MC, Cotter PD, Hill C, Ross RP. Fighting biofilms with lantibiotics and other groups of bacteriocins. npj Biofilms and Microbiomes. 2018;4(1):9.

12. Marković KG, Grujović MŽ, Koraćević MG, Nikodijević DD, Milutinović MG, Semedo-Lemsaddek T, et al. Colicins and Microcins Produced by Enterobacteriaceae: Characterization, Mode of Action, and Putative Applications. International Journal of Environmental Research and Public Health. 2022;19(18):11825.

13. De Silva PM, Bennett RJ, Kuhn L, Ngondo P, Debande L, Njamkepo E, et al. Escherichia coli killing by epidemiologically successful sublineages of Shigella sonnei is mediated by colicins. EBioMedicine. 2023;97.

14. Nedialkova LP, Denzler R, Koeppel MB, Diehl M, Ring D, Wille T, et al. Inflammation fuels colicin Ib-dependent competition of Salmonella serovar Typhimurium and E. coli in enterobacterial blooms. PLoS pathogens. 2014;10(1):e1003844.

15. Calcuttawala F, Pal A, Nath P, Kar R, Hazra D, Pal R. Structural and functional insights into colicin: a new paradigm in drug discovery. Archives of Microbiology. 2022;204(1):37.

16. Spriewald S, Glaser J, Beutler M, Koeppel MB, Stecher B. Reporters for single-cell analysis of colicin Ib expression in Salmonella enterica serovar Typhimurium. PLoS One. 2015;10(12):e0144647.

17. Isaacson R, Konisky J. Studies on the regulation of colicin Ib synthesis: I. Isolation of the Col Ib-P9 plasmid. Molecular and General Genetics MGG. 1974;132(3):215–21.

18. Askari N, Ghanbarpour R. Molecular investigation of the colicinogenic Escherichia coli strains that are capable of inhibiting E. coli O157: H7 in vitro. BMC veterinary research. 2019;15:1–8.

19. Nedialkova LP, Sidstedt M, Koeppel MB, Spriewald S, Ring D, Gerlach RG, et al. Temperate phages promote colicin-dependent fitness of S almonella enterica serovar T yphimurium. Environmental microbiology. 2016;18(5):1591–603.

20. Zhang Z, Du W, Wang M, Li Y, Su S, Wu T, et al. Contribution of the colicin receptor CirA to biofilm formation, antibotic resistance, and pathogenicity of Salmonella Enteritidis. Journal of basic microbiology. 2020;60(1):72–81.

21. Kong C, de Jong A, de Haan BJ, Kok J, de Vos P. Human milk oligosaccharides and non-digestible carbohydrates reduce pathogen adhesion to intestinal epithelial cells by decoy effects or by attenuating bacterial virulence. Food Research International. 2022;151.

22. Buchanan SK, Lukacik P, Grizot S, Ghirlando R, Ali MM, Barnard TJ, et al. Structure of colicin I receptor bound to the R-domain of colicin Ia: implications for protein import. The EMBO journal. 2007;26(10):2594–604.

23. Stecher B, Denzler R, Maier L, Bernet F, Sanders MJ, Pickard DJ, et al. Gut inflammation can boost horizontal gene transfer between pathogenic and commensal Enterobacteriaceae. Proceedings of the National Academy of Sciences. 2012;109(4):1269–74.

24. Feldgarden M, Riley MA. High levels of colicin resistance in Escherichia coli. Evolution. 1998;52(5):1270–6.

25. Davies JK, Reeves P. Genetics of resistance to colicins in Escherichia coli K-12: cross-resistance among colicins of group B. Journal of bacteriology. 1975;123(1):96–101.

26. Miller EA, Elnekave E, Flores-Figueroa C, Johnson A, Kearney A, Munoz-Aguayo J, et al. Emergence of a novel Salmonella enterica serotype reading clonal group is linked to its expansion in commercial Turkey production, resulting in unanticipated human illness in North America. MSphere. 2020;5(2):10.1128/msphere.00056-20.

27. Fornelos N, Browning DF, Butala M. The use and abuse of LexA by mobile genetic elements. Trends in microbiology. 2016;24(5):391–401.

28. Seo SW, Kim D, Latif H, O’Brien EJ, Szubin R, Palsson BO. Deciphering Fur transcriptional regulatory network highlights its complex role beyond iron metabolism in Escherichia coli. Nature communications. 2014;5(1):4910.

29. Drissi F, Buffet S, Raoult D, Merhej V. Common occurrence of antibacterial agents in human intestinal microbiota. Frontiers in Microbiology. 2015;6:441.

30. Datsenko KA, Wanner BL. One-step inactivation of chromosomal genes in Escherichia coli K-12 using PCR products. Proceedings of the National Academy of Sciences. 2000;97(12):6640–5.

31. Hoiseth SK, Stocker B. Aromatic-dependent Salmonella typhimurium are non-virulent and effective as live vaccines. Nature. 1981;291(5812):238-9.

32. Guzman L-M, Belin D, Carson MJ, Beckwith J. Tight regulation, modulation, and high-level expression by vectors containing the arabinose PBAD promoter. Journal of bacteriology. 1995;177(14):4121–30.

33. Chang AC, Cohen SN. Construction and characterization of amplifiable multicopy DNA cloning vehicles derived from the P15A cryptic miniplasmid. Journal of bacteriology. 1978;134(3):1141–56.

34. Zhou Z, Alikhan NF, Mohamed K, Fan Y, Agama Study G, Achtman M. The EnteroBase user’s guide, with case studies on *Salmonella* transmissions, *Yersinia* pestis phylogeny, and *Escherichia* core genomic diversity. Genome Res. 2020;30(1):138–52.

35. Zhou Z, Alikhan N-F, Sergeant MJ, Luhmann N, Vaz C, Francisco AP, et al. GrapeTree: visualization of core genomic relationships among 100,000 bacterial pathogens. Genome research. 2018;28(9):1395–404.

36. Ye J, McGinnis S, Madden TL. BLAST: improvements for better sequence analysis. Nucleic acids research. 2006;34(suppl_2):W6–W9.

37. Alikhan NF, Zhou Z, Sergeant MJ, Achtman M. A genomic overview of the population structure of *Salmonella*. PLoS Genet. 2018;14(4):e1007261.

38. Silva M, Machado MP, Silva DN, Rossi M, Moran-Gilad J, Santos S, et al. chewBBACA: A complete suite for gene-by-gene schema creation and strain identification. Microbial genomics. 2018;4(3):e000166.

39. Letunic I, Bork P. Interactive Tree of Life (iTOL) v6: recent updates to the phylogenetic tree display and annotation tool. Nucleic Acids Research. 2024:gkae268.

40. Schneider T, Hahn-Löbmann S, Stephan A, Schulz S, Giritch A, Naumann M, et al. Plant-made Salmonella bacteriocins salmocins for control of Salmonella pathovars. Scientific reports. 2018;8(1):1–10.

41. Cascales E, Buchanan SK, Duché D, Kleanthous C, Lloubes R, Postle K, et al. Colicin biology. Microbiology and molecular biology reviews. 2007;71(1):158–229.

42. Honeycutt JD, Wenner N, Li Y, Brewer SM, Massis LM, Brubaker SW, et al. Genetic variation in the MacAB-TolC efflux pump influences pathogenesis of invasive Salmonella isolates from Africa. PLoS Pathogens. 2020;16(8):e1008763.

43. Aljahdali NH, Khajanchi BK, Weston K, Deck J, Cox J, Singh R, et al. Genotypic and phenotypic characterization of incompatibility group fib positive salmonella enterica serovar typhimurium isolates from food animal sources. Genes. 2020;11(11):1307.

44. Upatissa S, Mitchell RJ. The “cins” of our fathers: rejuvenated interest in colicins to combat drug resistance. Journal of Microbiology. 2023;61(2):145–58.

45. Gilchrist JJ, MacLennan CA. Invasive nontyphoidal Salmonella disease in Africa. EcoSal Plus. 2019;8(2):10.1128/ecosalplus.ESP-0007-2018.

46. Suez J, Porwollik S, Dagan A, Marzel A, Schorr YI, Desai PT, et al. Virulence gene profiling and pathogenicity characterization of non-typhoidal Salmonella accounted for invasive disease in humans. PloS one. 2013;8(3):e58449.

47. Gal-Mor O, Boyle EC, Grassl GA. Same species, different diseases: how and why typhoidal and non-typhoidal Salmonella enterica serovars differ. Frontiers in microbiology. 2014;5:391.

48. Parsons BN, Humphrey S, Salisbury AM, Mikoleit J, Hinton JC, Gordon MA, et al. Invasive non-typhoidal Salmonella typhimurium ST313 are not host-restricted and have an invasive phenotype in experimentally infected chickens. PLoS neglected tropical diseases. 2013;7(10).

49. Kumwenda B, Canals R, Predeus AV, Zhu X, Kröger C, Pulford C, et al. Salmonella enterica serovar Typhimurium ST313 sublineage 2.2 has emerged in Malawi with a characteristic gene expression signature and a fitness advantage. Microlife. 2024:uqae005.

